# High efficiency genomic editing in Epstein-Barr virus-transformed lymphoblastoid B cells

**DOI:** 10.1101/379461

**Authors:** Andrew D. Johnston, Claudia A. Simões-Pires, Masako Suzuki, John M. Greally

## Abstract

While lymphoblastoid cell lines (LCLs) represent a valuable resource for population genetic studies, they are usually regarded as difficult for CRISPR-mediated genomic editing. It would be valuable to be able to take the results of their functional variant studies and test them in the same LCLs. We describe a protocol using a single-stranded donor oligonucleotide (ssODN) strategy for ‘scarless’ editing in LCLs. The protocol involves optimized transfection, flow cytometric sorting of transfected cells to single cells in multi-well plates and growth in conditioned, serum-rich medium, followed by characterization of the clones. Amplicon sequencing reveals the relative proportions of alleles with different editing events, with sequencing of DNA from clones showing the frequencies of events in individual cells. We find 12/60 (20%) of clones selected in this manner to have the desired ssODN-mediated recombination event. Long-range PCR of DNA at the edited locus and of RT-PCR products for the gene traversing the edited locus reveals 3/6 characterized clones (50%) to have large structural mutations of the region that are missed by sequencing just the edited site. The protocol does not require the use of lentiviruses or stable transfection, and makes LCLs a realistic cell type for consideration for CRISPR-mediated genomic targeting.

## INTRODUCTION

Lymphoblastoid cell lines (LCLs) have been collected for decades as a self-renewing cellular resource from different individuals, initially focusing on multi-generational families to facilitate linkage mapping (1). In recent years, they have been collected as a resource for population genetics studies, by the HapMap Consortium (2), the GEUVADIS Project (3), and the 1000 Genomes Project (4), among others. Repositories such as the Coriell Institute and the American Type Culture Collection have between them banked tens of thousands of LCLs, indicating the scale at which these resources are being generated. LCLs continue to be successfully exploited as a model system for understanding how DNA sequence variability leads to individual differences in transcriptional regulatory variation. LCL panels have also proven useful in pharmacogenomic studies (5). Furthermore, individual LCLs have served as reference cell lines for a number of purposes. The 17 member, 3 generation CEPH pedigree 1463 has been the target of Illumina’s Platinum Genomes sequencing, while the mother in the second generation of this family is the GM12878 cell line, a tier 1 ENCODE cell line (6) that has been used in many novel assays, including some of ours (7), assays studying chromatin looping (8), and the Genome in a Bottle initiative (9). The position of LCLs in general and some lines in particular at the center of many modern molecular genomic discoveries is clear.

However, the discovery of functional variants in LCLs is rarely followed by genomic editing in the same cell type (10) unless lentiviral transfection is used to express Cas9 robustly (11). If the locus containing the functional variant is very LCL-specific in its activity, it may be difficult to test the effect of DNA sequence polymorphism at the locus in a surrogate cell type. Furthermore, if LCLs are to be used in CRISPR screens of the whole genome (12), an effective protocol for its use in this cell type is going to be needed. The most successful reported use to date of CRISPR in LCLs does not describe the efficiency of their strategy, making it difficult to replicate (10). We therefore sought to test how efficiently homology-directed recombination (HDR) using a single-stranded donor oligonucleotide (ssODN) could be used for ‘scarless’, CRISPR-mediated editing in LCLs.

## RESULTS

### CRISPR/Cas9 editing yields a high proportion of successfully-edited clones

Our protocol overview is shown in **Figure 1**. The oligonucleotides used for this study are provide in **Table 1**. The protocol details are in the **Methods** section. We started with 4 × 10^6^ LCL cells and 33.3 μg of plasmid for transfection, observing this to be accompanied by ~40% cell death. Overnight incubation allowed the GFP to be expressed in transfected cells, allowing fluorescence-activated cell sorting (FACS) to deliver individual cells into 96 well plates. We plated over 2,000 individual cells and allowed the cells to grow in conditioned, FBS-rich medium for 2 weeks. Almost one-quarter of the plated individual cells were growing at this point. We re-plated a total of 480 cells, and chose the 60 fastest-growing cells colonies for screening. We performed PCR to amplify the targeted region, and then digested the PCR product from each clone with the Styl restriction endonuclease, as its recognition site, present in the unedited cell line, should be disrupted by any genomic editing event (**Figure 2a**). This revealed 55 of the 60 clones to have had some sort of editing event at the locus, prompting the sequencing of these amplicons, and revealing that 12 of the 55 had apparent homozygosity for the desired homology-directed recombination (HDR) event.

**Figure 1:**
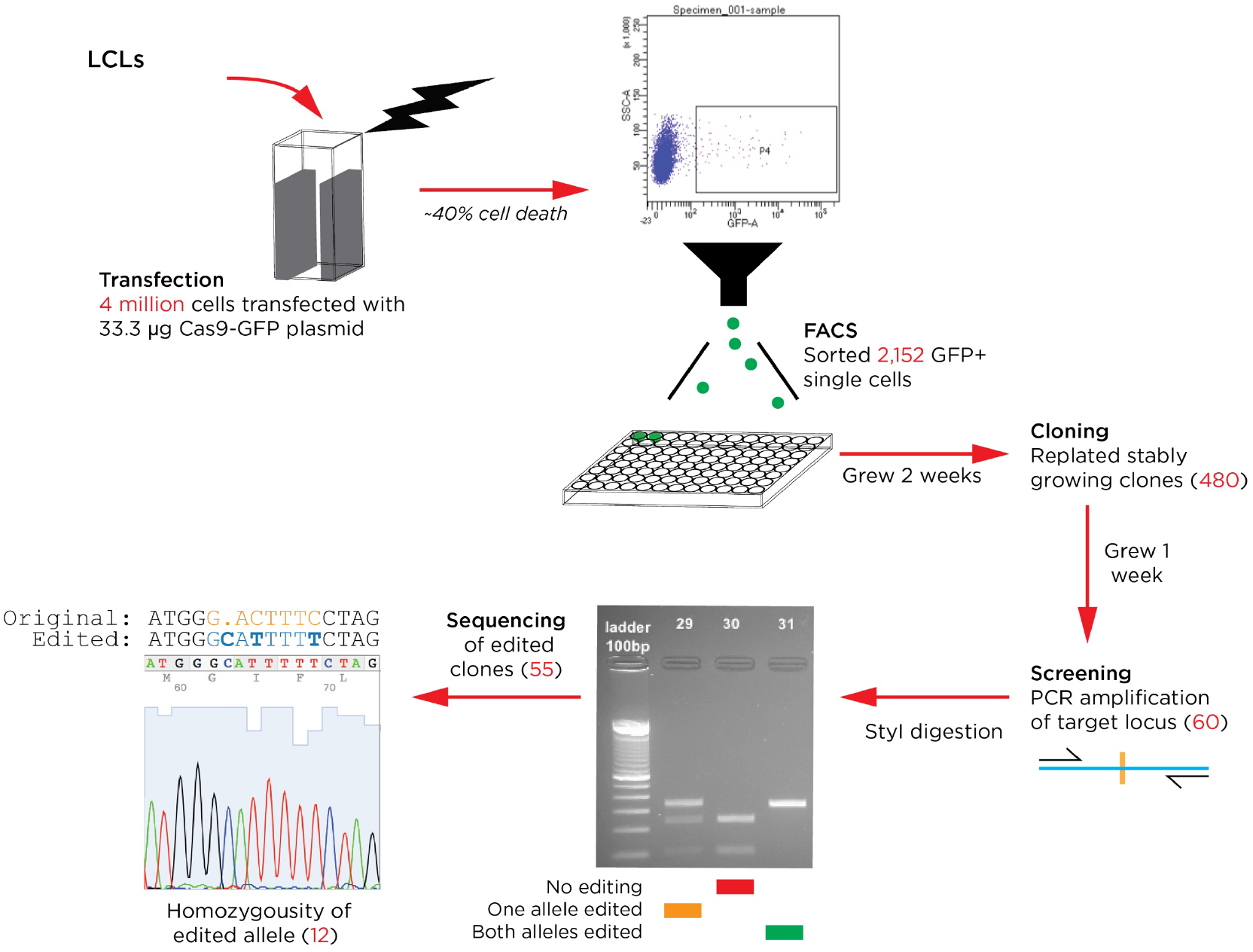
Overview of the LCL editing protocol. The numbers of cells at each stage are shown. Substantial cell death occurs following transfection, but the cloning steps were relatively efficient, and the rate of recovery of edited clones was high.

**Figure 2:**
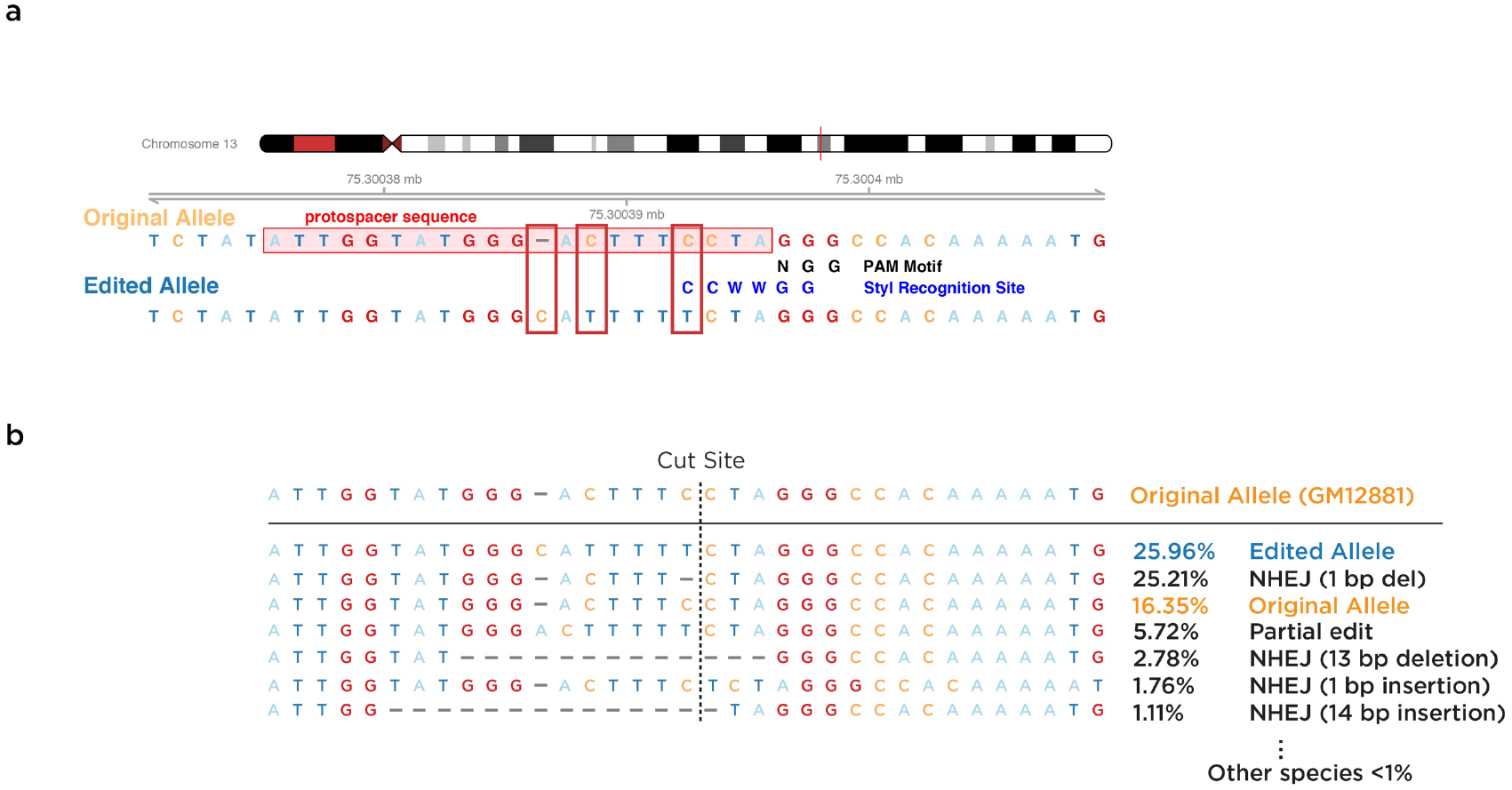
(a) The characteristics of the edited locus are shown. The protospacer sequence, to which the guide RNA binds, is shown to be at the same site as is being targeted for editing by the ssODN, which then prevents the guide RNA from binding to cause further edits. The location of a StyI recognition motif present in the unedited DNA is shown, demonstrating how restriction enzyme digestion with StyI for this particular editing event can be used in screening for editing events. In (b) we show the results of amplicon sequencing, and the relative frequencies of each type of editing event. The desired editing event is the most common event, followed by non-homologous end joining (NHEJ) with deletion of the single nucleotide immediately at the cut site.

**Table 1:**
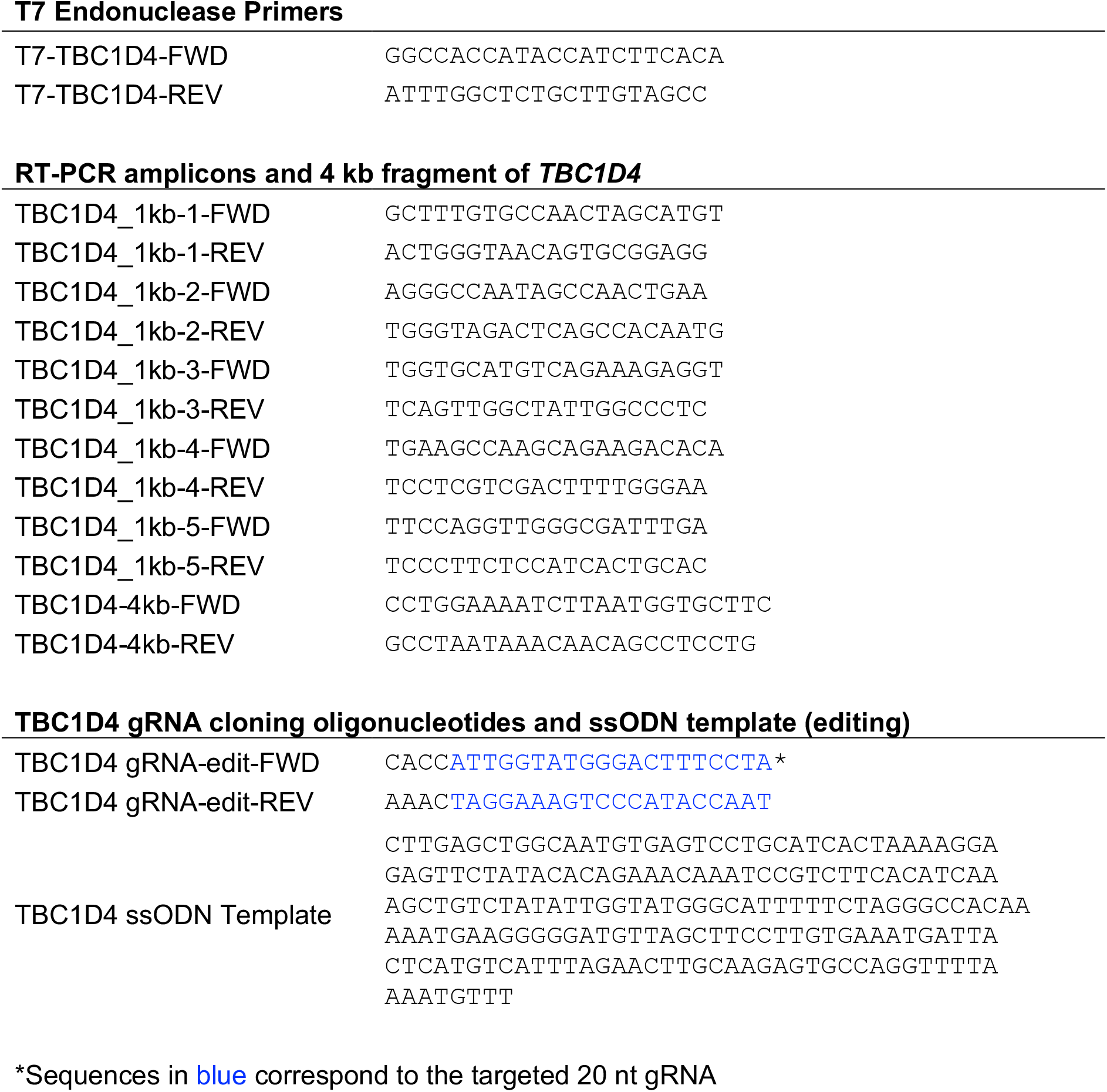
Oligonucleotides used in project.

### Amplicon-seq reveals the distribution of different types of editing events

To gain a more comprehensive insight into the editing occurring in the cells, we performed amplicon sequencing of the pool of cells prior to the cloning step. We show the results of analysis of these data using CRISPResso (13) in **Figure 2b**. The amplicon PCR primers were designed to flank and avoid overlapping the ssODN sequence. The most common edit was the desired HDR (25.96%), followed by the 1 bp deletion immediately upstream of the cut site representing non-homologous end-joining (NHEJ, 25.21%), with only 16.35% of the sequences representing unedited alleles. The remaining approximately one-third of alleles represented a wide range of events.

In **Figure 3**, we show the results of another experiment testing the effect of using the singlestranded oligonucleotide (ssODN) template. In **Figure 3a**, we show that the proportion of events with HDR without using the ssODN was zero, but rose to 0.233 when the ssODN was used. In **Figure 3b**, we separate events involving just HDR from those which also involve other types of NHEJ-mediated repair. By definition, the desired insertion and substitutions occur at 100% frequency in the HDR-only alleles, but when we add back the NHEJ events, we see that they are most frequent immediately adjacent to the cut site, and fall off in frequency even within the first 10 bp from the cut site.

**Figure 3:**
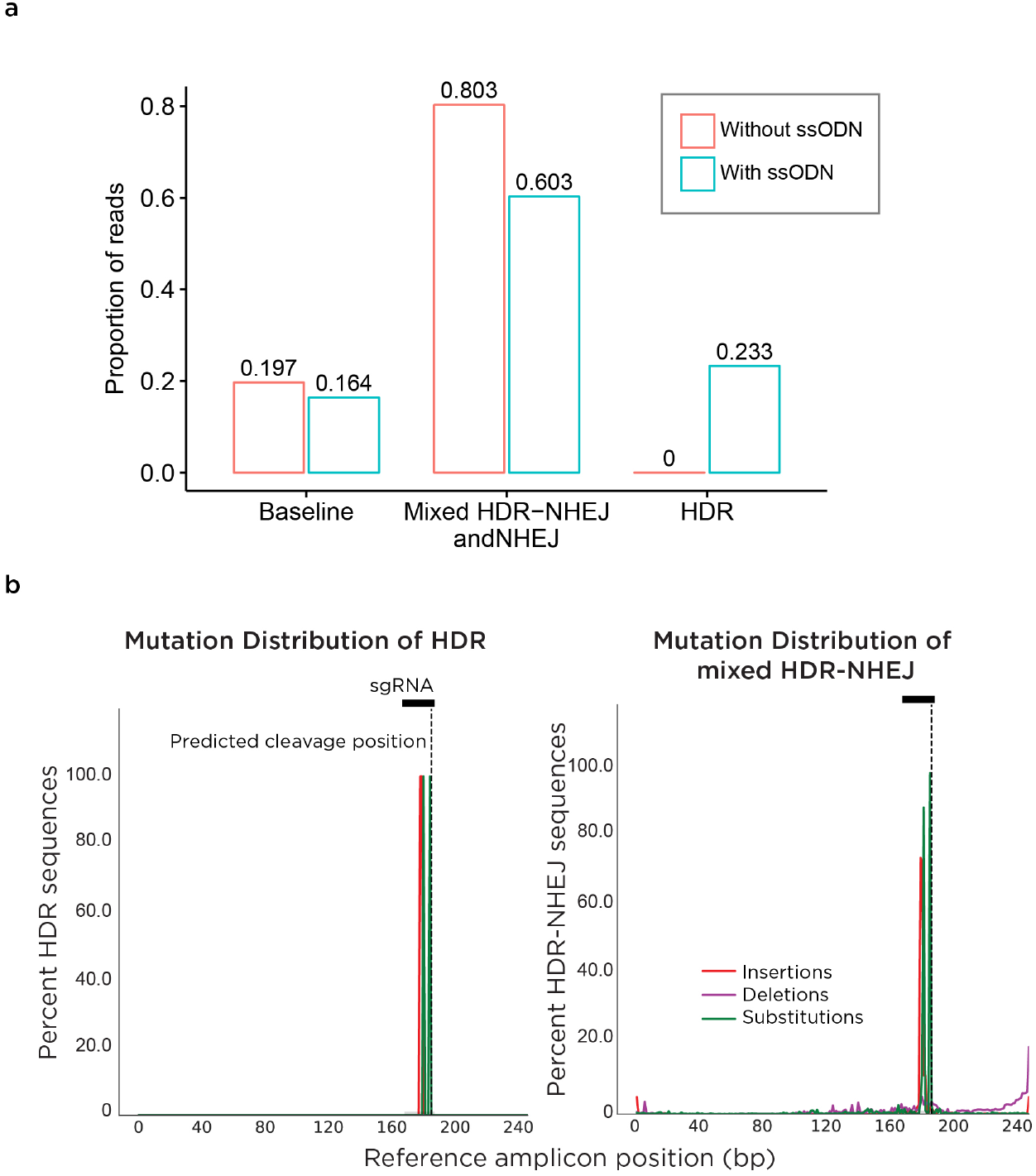
(a) The use of the ssODN is responsible for all HDR events, bringing its frequency from zero to over 23%. In (b) we show the three events in the HDR alleles, allowing us to appreciate what is illustrated on the right, that NHEJ events occur more frequently closer to the cut site and diminish in frequency dramatically over <10 bp.

### Testing for large rearrangements at the targeted locus

As it has recently been described that CRISPR-mediated genomic editing can induce large structural rearrangements that would be missed when testing just the immediate <200 bp around the target site (14), we performed two types of screening to test for these events. One strategy was to perform long-range PCR in the 4 kb surrounding the locus, making sure neither primer was near the cut site or overlapped the ssODN sequence. As the targeted locus is within an intron of a gene that is transcribed over almost 200 kb of the genome, we also designed RT-PCR primers, each amplifying ~1 kb fragments, spanning multiple exons and thus a larger region flanking the CRISPR target. We show the characteristics of the 5 cell samples tested in **Figure 4a** and the design of these primers in **Figure 4b**. In **Figure 4c** we show that two clones (one with a 14 bp deletion, the other with a combined edit and indel) showed PCR products of <4 kb from the DNA amplification assay. The RT-PCR amplicons were compared by densitometry and normalised to the signal obtained for the amplification from the 3’ UTR. We show these results in **Figure 4d**. While there are differences overall in amplification efficiency for each primer pair, we observe the 7 bp deletion and the 14 bp deletion clones individually moving outside the range of values of the unedited, original and other clones tested. We show in **Figure 4b** where this indicates there to be larger-scale rearrangements in the clones tested, confirming the need for caution in performing these editing experiments (14), and demonstrating that the apparent homozygosity observed for the 7 bp deletion clone is instead likely to be due to hemizygosity at this locus.

**Figure 4:**
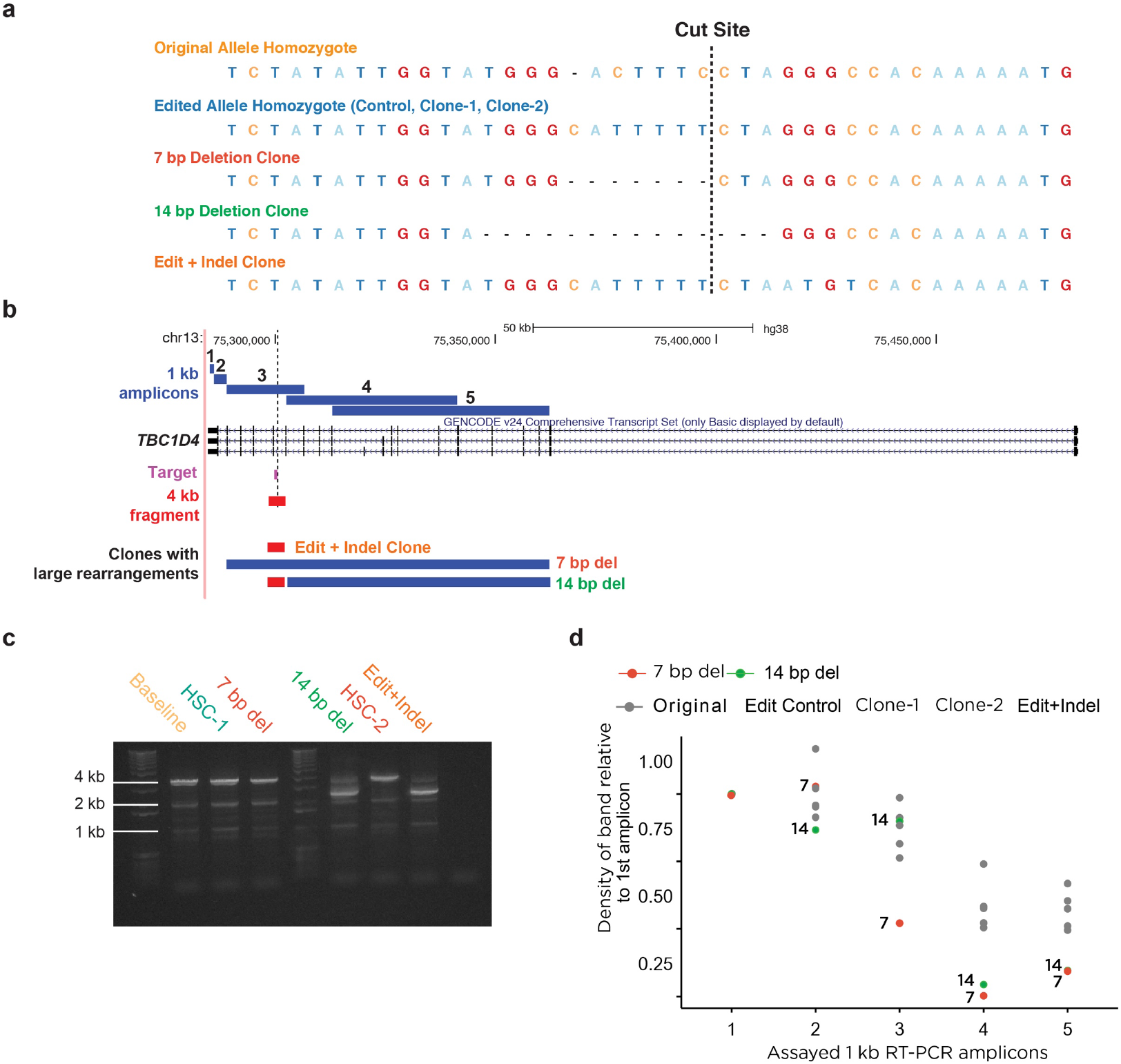
In (a) we show the unedited sequence and the editing events at the target site in 4 different clones. In (b) we show the location of the 4 kb PCR of the DNA flanking the edited region (red), and the five sets of RT-PCR primers amplifying 1 kb fragments of the *TBC1D4* mRNA. In (c) we show the results of the 4 kb DNA PCR, showing two clones (14 bp deletion and the edit/indel clone) to have <4 kb amplicons, indicating that these have deletions within the amplified region. In (d) we show the densitometry of the RT-PCR products normalized to the 3’ UTR signal. The 7 and 14 denote the 7 bp and 14 bp deletion clones, respectively. The range of values shows the 7 bp and 14 bp deletion clones to be outliers, indicating differing sized deletions in each clone. The likely locations of deletions are represented at the bottom of panel (b).

## DISCUSSION

As has been demonstrated in a prior report, LCLs can be edited using CRISPR and HDR to modify a locus to change from one to another human single nucleotide variant at a locus regulating local gene expression (10). We take this observation further to reveal the critical steps in the protocol, not just the points of greatest inefficiencies, but also some surprising efficiencies.

We note that the transfection rate is highly sensitive to the number of cells used, requiring that very accurate cell counting be performed. We confirm some previous experimental design optimizations for improved HDR, including the design of ssODNs that do not hybridize with gRNAs or the template strand of transcribed genes, and the use of a double-stranded nuclease instead of nickase (15). By targeting the HDR-directed mutations to within the guide RNA binding site, we reduce the re-cutting of the site by CRISPR to enhance scarless editing (16). Furthermore, we confirm that cuts close to the target edit site are more efficient (15, 17).

Our study did not encompass testing for off-target effects. We attempted to guard against this by using a high-fidelty espCas9(1.1) that is created through mutations that affect its propensity to cleave when there are mismatches between the gRNa and the protospacer (18). The group who designed this Cas9 did not observe any edits at common off-target sites when these were were assessed (18), which should apply in LCLs to the same extent as other cells.

There are some simple guidelines worth following when editing LCLs. The protocol outline shown in **Figure 1** provides some guidance about cell numbers at each step. We had excessive cell loss during culture because the wells at the outside of the multi-well plate evaporated more quickly. A trick for the future would be to keep those wells filled with water and use only the internal wells for cell culture. Plasmids need to be highly concentrated to avoid diluting nucleofection reagents. We anticipate that the efficiency of the protocol will improve markedly over time. With the introduction of ribonucleoprotein reagents for guide RNAs and Cas9, the use of plasmids will possibly diminish. All of the techniques we used are potentially automated by liquid handling/cell culture systems, raising the possibility that LCL editing may be amenable to scaling to many cell lines at a time, potentially introducing the same variant in multiple genetic backgrounds to create a system for studying phenomena like epistasis, lending further value to this research workhorse cell type.

## METHODS

### Cell line culture

The lymphoblastoid cell line (LCL) derived from a child within the CEPH Pedigree 1463 (GM12881) was purchased from the Coriell Institute and cultured in RPMI 1640 medium, supplemented with 15% fetal bovine serum (FBS, Benchmark), 100 IU/ml penicillin, and 100 μg/ml streptomycin (Life Technologies). Cells were kept in suspension in tissue culture flasks (NUNC, Thermo Scientific) at 37 °C in a 5% CO_2_ incubator and maintained between 2 × 10^5^ and 8 × 10^5^ cells/ml.

### Plasmids and ssODN template

Plasmid pCAG-eCas9-GFP-U6-gRNA was a gift from Jizhong Zou (Addgene plasmid #79145) and was used in combination with a single-stranded oligonucleotide (ssODN) template (**Table S1**) for editing purposes. Plasmid pmaxGFP (Lonza) was used as GFP positive control vector. gRNAs were designed using the CRISPOR web interface (19). To insert the desired gRNA sequence into the CRISPR/Cas9 vector, reverse complement oligonucleotides containing the 20-nt gRNA target sequence (**Table S1**) were annealed, 5’-phosphorylated and ligated into the linearized vector.

### CRISPR/Cas9 editing and sorting

For transfection, cells were passaged at 3.5×10^5^ 48 hours and 24 hours before transfection. A total of 4 × 10^6^ GM12881 cells were transfected with 33.3 μg of CRISPR/Cas9 plasmid and 0.4 nmol of ssODN template. GFP control cells received 2 μg of GFP plasmid, while negative control cells received transfection reagents only. Transfections were conducted with the Cell Line Nucleofactor Kit V (Lonza) according to the manufacturer’s instructions. After transfection, cells were suspended in medium and incubated overnight under normal cell culture conditions, then replaced with fresh medium. GFP-positive cells were sorted after 48 hours following the transfection. The cells were pelleted, washed twice, and suspended in sorting buffer (Hank’s balanced salt solution buffer supplemented with 1% FBS, 100 units/mL penicillin, and 100 μg/mL streptomycin). Cell suspensions were submitted to cell analysis and sorting in a FACSAria II cytometer (BD Biosciences). FACS data were analyzed using FACSDiva software (Becton Dickinson) with gating of single cells using FSC/W and SSC/W, and gating of GFP-positive cells.

### Clone isolation upon genomic editing

Single GFP-positive cells were sorted 48 hours after transfection into individual wells of a 96-well plate, containing a mixture of fresh and conditioned medium (1:1), in which the FBS concentration was increased to 20%. The 96-well plate was incubated for two weeks under cell culture conditions, then the clones exhibiting robust growth were transferred to a new 96-well plate with additional conditioned medium. Given the high concentration of FBS and the long culture times, a precipitate may form, which can easily be mistaken for contamination. Driven by evaporation, we recommend plating water in the peripheral wells of a 96-well plate to avoid disruption of cell growth by crystal formation. Conditioned medium was obtained from GM12881 cells, and cultured in 20% FBS RPMI 1640 for 24 hours. The medium was removed without disturbing cells at the bottom of the flask, centrifuged at 2,000 rpm, and the supernatant was filtered through a 0.3 micron sterile filter prior to use. When subsequent T7EI assays were to be performed, cells were sorted directly into QuickExtract DNA extraction solution (Epicentre).

### T7 endonuclease I assay (T7EI)

To verify editing in clones, in which an intronic enhancer of *TBC1D4* was targeted, genomic DNA was isolated from transfected and control cell pellets using QuickExtract DNA Extraction Solution (Epicentre) according to manufacturer’s instructions. DNA was then concentrated by ethanol precipitation. The 1 kb region containing the gRNA targeted region was amplified with forward and reverse primers (**Table S1**) using the Q5 Hot Start High-Fidelity 2X Master Mix (NEB) according to the manufacturer’s protocol with 100 ng of the purified total cellular DNA in a 50 μl reaction. Amplification products were isolated using the DNA Clean and Concentrator-5 kit (Zymo Research). PCR product (50 ng) was denatured and re-annealed in a final volume of 13 μl in 1X NEBuffer2 (NEB) using a thermocycler with the following protocol: 95°C, 5 minutes; 95➔85°C at −2°C/second; 85➔25°C at −0.1 °C/second; hold at 4 °C. Hybridized PCR products were treated with 10 U of T7E1 enzyme (NEB) at 37 °C for 60 minutes in a reaction volume of 20 μL. The reaction was stopped with 2 μL of 0.25 M EDTA, and subsequently analyzed on a 1.5% agarose gel.

### Amplicon-seq generation and data analysis

Cell lines generated by CRISPR/Cas9 editing at a locus with or without a repair template were assessed by amplicon-seq. A suspension of 10^4^ cells sorted in QuickExtract DNA extraction solution (Epicentre) was used for DNA extraction according to manufacturer’s instructions. DNA was extracted by vortexing the cell suspension for 15 seconds, followed by incubation at 65°C for 6 minutes, an additional 15 second vortexing, and a final 2 minute incubation at 98°C. DNA was then concentrated by ethanol precipitation and submitted to an initial PCR with locus-specific forward and reverse primers with portions of the Illumina TruSeq adapters on their 5’ ends. PCR products were purified with DNA Clean and Concentrator-5 kit (Zymo), and a second round of PCR was performed on purified DNA with primers containing the remaining Illumina adapter along with a custom 6-nt index on the reverse primer. The amplicon libraries were then purified by gel extraction and sequencing using Illumina MiSeq technology, 250 bp single end sequencing. The resulting data were analysed using CRISPResso (13).

### RT-PCR of the *TBC1D4* 1 kb amplicon

Cell pellets were treated with QIAzol lysis reagent (Qiagen) and total RNA was isolated using the miRNAeasy kit (Qiagen) combined with on column DNAse (Qiagen) treatment according to manufacturer’s instructions. Synthesis of cDNA was performed with total RNA and SuperScript III First-Strand Synthesis System for RT-PCR (Life technologies) using oligo(dT)20 as primers. Subsequent PCR was conducted with primers designed using the NCBI Primer-BLAST web interface (20) (**Table S1**) and the Q5 hot start high fidelity polymerase master mix, according to the manufacturer’s protocol. Samples were then analyzed on a 2% agarose gel.

## ACCESSION NUMBERS

All genome sequencing data are available from the NCBI Gene Expression Omnibus database under accession number GSE117576 (https://www.ncbi.nlm.nih.gov/geo/query/acc.cgi?acc=GSE117576).

## SUPPLEMENTARY DATA

Supplementary Data are available at NAR online. One Supplementary Table is provided with oligonucleotide descriptions needed to replicate this study.

## ACKNOWLEDGEMENT

The authors thank the following Einstein core facilities for their expertise: the Epigenomics Shared Facility, the Flow Cytometry Core Facility and the Genomics Core Facility.

